# The nematode effector Mj-NEROSs interacts with ISP influencing plastid ROS production to suppress plant immunity

**DOI:** 10.1101/2022.10.24.513376

**Authors:** Boris Stojilković, Hui Xiang, Yujin Chen, Lander Bauters, Hans Van de Put, Kathy Steppe, Jinling Liao, Janice de Almeida Engler, Godelieve Gheysen

**Affiliations:** Department of Biotechnology, Ghent University, Proeftuinstraat 86, Ghent 9000, Belgium; Laboratory of Plant Nematology, South China Agricultural University, Guangzhou 510642, China; Guangdong Province Key Laboratory of Microbial Signals and Disease Control, South China Agricultural University, Guangzhou 510642, China; Guangdong Vocational College of Ecological Engineering, Guangzhou 510520, China; Laboratory of Plant Ecology, Department of Plants and Crops, Faculty of Bioscience Engineering, Ghent University, Coupure links 653, 9000, Gent, Belgium; INRAE, Université Côte d’Azur, CNRS, ISA, Sophia Antipolis, France

**Author notes:** **Correspondence** and requests for materials should be addressed to Godelieve Gheysen. Co-second authors, have contributed equally (Hui Xiang, Yujin Chen).

**Keywords:** Plant defence, ROS, root plastids, effector, ISP, protein-protein interaction

## Abstract

Root-knot nematodes are an important group of plant pathogens that mainly infect plant roots. They establish a feeding site in the host upon infection while secreting hundreds of effectors. These effector proteins are crucial for successful pathogen propagation. Although many effectors have been described, their targets and molecular mode of action are still unknown. Here we report the analysis of the RKN effector, Mj-NEROSs (***M****eloidogyne* ***j****avanica* **n**ematode **e**ffector **ROS s**uppressor), which emerges to have an essential role in suppressing host immunity by inhibiting INF1-induced cell death and reducing callose deposition. Secreted from the subventral esophageal glands to giant cells, Mj-NEROSs localizes in plastids where it interacts with **ISP**, interfering with the electron transport rate and ROS production. Moreover, our transcriptome analysis shows the downregulation of ROS-related genes upon Mj-NEROSs expression. We propose that Mj-NEROSs manipulates root plastids leading to transcriptional changes, lowering ROS production, and suppressing host immunity.

## Introduction

Root-knot nematodes (*Meloidogyne spp*.) are obligate plant pathogens that infect many important plant crops ^1^. After the invasion of the plant root near the tip, root-knot nematodes (RKNs) juveniles migrate to the vascular cylinder and become sedentary. Selected root cells from the vascular parenchyma are hijacked by the nematodes and then undergo numerous rounds of karyokinesis without cytokinesis, resulting in multinucleate giant-feeding cells. Besides multiple nuclei and vacuole fragmentation, giant cells are characterized by cytoskeleton rearrangements and plastid differentiation into chloroplast-like organelles ^2^ accompanied by a remarkable induction of genes involved in chloroplast biogenesis ^3^. Several multinucleate giant cells are part of a feeding site that serves as the primary source of nutrients for this root pathogen’s development and maturation ^4^. Keeping the feeding structures alive in their host is crucial for the nematodes’ survival, and one essential mechanism in doing so is the suppression of the plant defense system. Root-knot nematodes can manipulate plant cells by secreting various molecules named effectors ^5–7^. Effectors can target a range of host structures, including the nucleus, plastids, cell wall, and molecular machinery involved in fundamental processes ^8–10^, such as the plant’s immunity ^11–13^.

On the other side, plants have evolved different strategies to combat pathogen infection. The first line of defense is based on recognizing pathogen-associated molecular patterns (PAMPs) and damage-associated molecular patterns (DAMPs) produced by wounded plant tissue ^14–16^. Recognition of these two groups of molecular patterns will activate PAMP-triggered immunity (PTI) ^15^. Pathogen effectors can suppress the PTI ^17^ but in case plants recognize the secreted effectors, the second layer, ETI (effector-triggered immunity), will be activated. ETI is a more rapid immune response, often associated with a localized hypersensitive response (HR) at the site of infection ^18^. Plant defense strategies include the production of reactive oxygen species (ROS), callose deposition, gene expression modification, and plastid deformation ^19,20^.

ROS appear to act as a critical component of the defense system ^21–23^. They mediate diverse plant defense responses through various signaling pathways, including protein phosphorylation-induced signaling cascades and pathways involving chemical signals (e.g., nitric oxide) or plant hormones (e.g., salicylic acid) ^24,25^. ROS represent a plant weapon in cross-kingdom interactions; either being directly toxic to a pathogen or as a cellular messenger that dictates inter-organelle communication and influences gene expression ^23,26^. The ROS production in plants is mainly localized in the chloroplasts, mitochondria, peroxisomes, and apoplast ^22,23,27,28^. Under unfavorable circumstances, especially in interacting with pathogens, plants generate ROS to regulate defense-modulating programmed cell death and hormonal production ^29–31^. Even though low levels of ROS are needed for several biological processes ^32^, high levels can lead to damage of the plant’s DNA and, consequently, cell death ^29,33^. Not surprisingly, many molecules involved in the ROS production cascade serve as an effector’s target in the constant fight of the pathogen to remain hidden from the plant immunity and to counteract the damaging action of ROS ^34,35^. Although this is previously described for many bacterial and fungal effectors ^36–39^, the precise mode of action of RKN effectors and their targets are still largely unknown ^8,9^. Only one RKN effector was shown to target plastid components and to suppress the oxidative burst in leaves ^40^.

Here, we report the characterization of the *M. javanica* putative effector, Mj-NEROSs, using an integrative approach previously used in the field ^41^. Mj-NEROSs is a homolog of Mi-4D01 (also referred to as Mi-Msp3^42^), a potential RKN effector that was firstly reported by Huang et al., (2003)^5^ and shown to be expressed in the subventral gland of *M. incognita*. The expression studies on Mi-4D01 showed maximum expression in the egg stage, subsequently declining in J2s and adult females ^42^. By yeast-two hybrid screening and confocal microscopy, we show that Mj-NEROSs localizes in chloroplasts, interacting with the plant’s Iron-Sulfur Proteins (ISP). ISP is a component of the cytochrome b6-f complex whose primary role is to mediate electron transfer in photosynthesis ^43–45^. In addition, this protein plays an essential role in plant immunity, and it has been previously described as a target of several fungal effectors ^36,46^. We deduce here that the binding of Mj-NEROSs to ISP affects the function of the Cyt b6/f complex in the electron transport chain and impairs ROS accumulation leading to changes in gene expression, resulting in the suppression of plant immunity.

## Results

### About NEROSs

To better understand the molecular mechanisms of the *M. javanica*– *Solanum lycopersicum* interaction, we investigated most of the predicted *M. incognita*/*M. javanica* effectors ^5,47–49^ for their potential localization. We found a substantial number (ninety) of effectors with potential chloroplast localization signal (**Supplementary Table 1**). One of them, *Meloidogyne incognita* predicted esophageal gland cell secretory protein 3 (AF531162) ^5^, was shown to have a significant impact on the life cycle completion in an RNAi study ^42^. We decided to investigate the function and mode of action of its closest *M. javanica* homolog that showed 96% identity on protein level (**Supplementary Fig. 1a**). Here, we aim to confirm its role in nematode parasitism and unveil the interplay between this RKN effector and the host plant. Based on our observations below, this protein will be referred to as Mj-NEROSs (***M****eloidogyne* ***j****avanica* **n**ematode **e**ffector **ROS s**uppressor). The predicted coding sequence of Mj-NEROSs encodes a protein of 174 amino acids with a calculated molecular size of 18.7 kDa. The N-terminal 24-amino acids are predicted to be a signal peptide responsible for the secretion of the effector protein to the plant cell. The version of the protein without this signal peptide is referred to as NEROSs.

### *Mj-NEROSs* is highly expressed in later parasitic stages

To determine the temporal resolution of *Mj-NEROSs*, its transcriptional levels were investigated at several developmental stages by quantitative RT-PCR (qRT-PCR). The expression of *Mj-NEROSs* was low in the early stages of the infection. However, this changed during the later time points (18 DPI (days post-infection) and 21 DPI), where the expression increased and reached its maximum. Finally, the mRNA levels of *Mj-NEROSs* decreased to its lowest levels in adult females (30 DPI) (**Fig. 1a**).

**Fig. 1.**
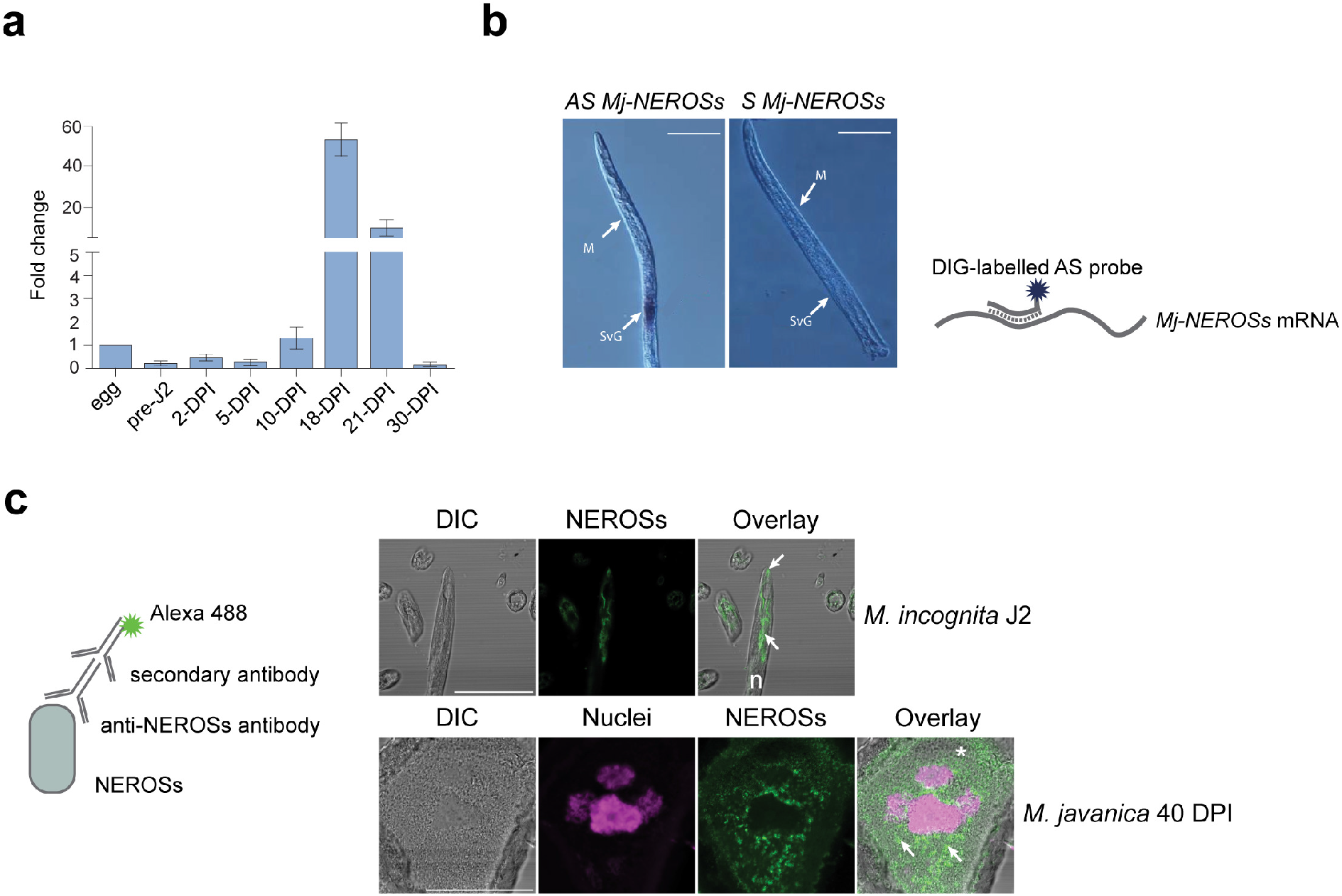
Temporal expression of *Mj-NEROSs* by *in-situ* hybridization and protein localization. **a** Relative expression levels of *Mj-NEROSs* in *Solanum lycopersicum* infected roots at different time points after inoculation (days post inoculation; DPI) assayed by qRT-PCR; *Mj-β-actin* represents a reference gene. Mean and standard deviations were calculated from three biological replicates, each containing three technical replicates. **b** *In-situ* hybridization (ISH) localized mRNA of *Mj-NEROSs* in the subventral gland of stage 2 juvenile (J2) of *M. javanica*. M, median bulb; SvG, subventral gland; AS, anti-sense; S, sense. **c** The upper panel illustrates the immunolocalization of NEROSs in SvG and secreted (white arrows) by pre-parasitic *M. incognita* J2. The lower panel, shows NEROSs fluorescence in giant cells foci, likely secreted by *M. javanica* 40 DPI in tomato roots. n, nematode; asterisk, giant cell; NEROSs, green fluorescence and white arrows, DAPI stained nuclei, magenta color. Scale bars 50 μm.

These findings implicate that Mj-NEROSs is present all through the parasitic cycle of *M. javanica*. However, it most likely plays a more significant role during the mid-later parasitic stages.

### NEROSs is expressed in the subventral glands and secreted in giant cells where it shows localization in giant cell foci

To investigate the spatio-temporal expression of *Mj-NEROSs* and to examine its protein location in the plant host during nematode parasitism, *in situ* hybridization (ISH) and immunolocalization (IL) were performed respectively. A substantial accumulation of *Mj-NEROSs* transcripts was observed in the subventral esophageal glands of pre-parasitic J2s (**Fig. 1b**), and no signal was observed during hybridization with the sense single-strand DNA used as a negative control (**Fig. 1b**). Immunolocalization results are in alignment with ISH showing that NEROSs protein is produced in the subventral esophageal glands of pre-parasitic juvenile nematodes before secretion. In addition, NEROSs fluorescence was observed at the nematode stylet (**Fig. 1c; upper panel**). Analysis of different time points after infection illustrated a visible fluorescence at 21 and 40 DPI (**Supplementary Fig. 1b**). The NEROSs fluorescence signal was observed specifically in the giant cell cytoplasm, and this pattern was similar for both analyzed nematodes (*M. javanica* and *M. incognita*) (**Supplementary Fig. 1b and Fig. 1c**). At 40 DPI we observed specific localization of NEROSs in giant cells. The signal was absent from nuclei and present in giant cell foci in the cytoplasm (**Supplementary Fig. 1b middle panel and Fig. 1c, lower panel**).

Overall, the data illustrated that Mj-NEROSs is expressed in the nematode subventral esophageal glands and secreted in giant cells at middle to middle-later stages of parasitism, where it localizes explicitly in giant cell cytoplasmic foci.

### NEROSs interacts with ISP (Y2H)

To understand the role that Mj-NEROSs plays in host plants, we first screened a yeast two-hybrid library of *Arabidopsis thaliana* roots (infected with nematodes) using NEROSs as bait. We found an interacting partner **(Supplementary Fig. 2a)** of NEROSs being an Iron-Sulfur Protein (AtISP), the component of the cytochrome b6/f complex involved in connecting photosystem II (**PSII**) to Photosystem I (**PSI**) in the electron transport chain across the chloroplast’s thylakoid membrane ^43,44^. To test whether this interaction was conserved among different host plants, we cloned the full-length coding sequence of ISP from *S. lycopersicum* Sl-ISP, here named ISP and *N. benthamiana* NbISP and screened for interaction with NEROSs in pairwise yeast two-hybrid. The interaction was found for the three species, being the strongest between NEROSs and ISP from tomato **(Fig. 2a and Supplementary Fig. 2a)**. Therefore, we decided to focus on unveiling the role of this interaction in *S. lycopersicum*.

**Fig. 2.**
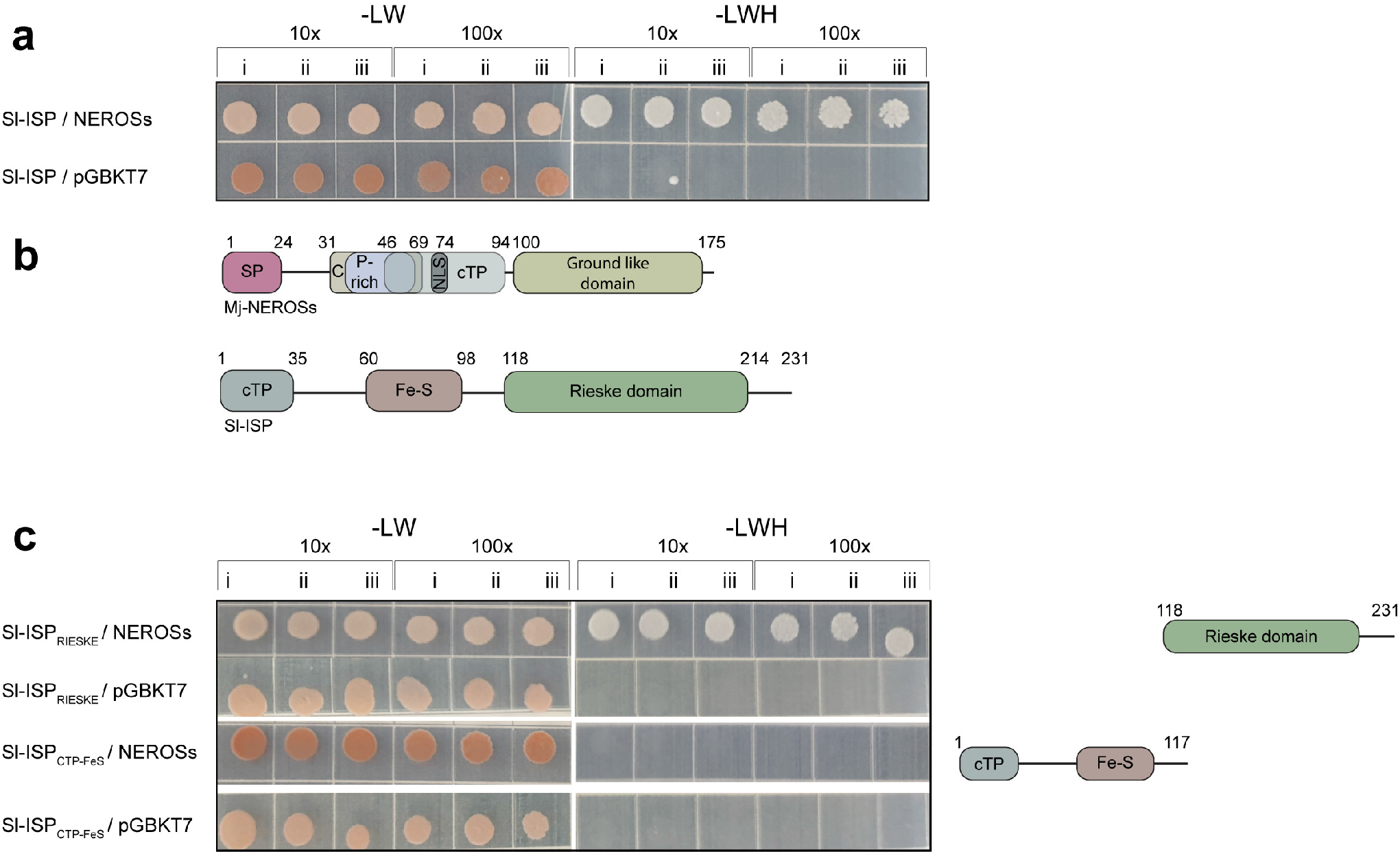
Domain organization and binding partner ISP. **a** Detection of the interaction between NEROSs and Sl-ISP by yeast two-hybrid assay. Yeast cells of the PJ69-4A strain transformed with the pGBKT7_*NEROSs* and pGADT7_*Sl-ISP* constructs; cells spotted on SD (Synthetic Defined) medium lacking LW (leucine and tryptophan) or LWH (lacking leucine, tryptophan, and histidine). Dilutions are indicated on top; i, ii, and iii indicate three different biological replicates. **b** Schematics of the Mj-NEROSs and SI-ISP domain organization. Top: the predicted signal peptide of Mj-NEROSs is followed by cysteine-rich (C) and proline-rich (P-rich) domains; chloroplast transport peptide (cTP) is followed by the C-terminal ground-like domain. Bottom: SI-ISP protein consisting of a chloroplast transport peptide (cTP) followed by CytB6 Fe-S and Rieske domains. Black lines represent disordered regions. **c** Determining the interacting parts of NEROSs and Sl-ISP by yeast two-hybrid assay. Yeast transformants expressing the truncated constructs of NEROSs and Sl-ISP domains were assayed for growth on SD-LW or SD-LWH. SI-ISP_RIESKE_ represents a version only with the Rieske domain; Sl-ISP_cTP-FeS_ contains the N-terminal region of Sl-ISP and the Fe-S domain. pGBKT7 is an empty vector. Schematics of the tested truncated protein versions are represented on the right; dilutions are indicated on top; i, ii, and iii indicate three different biological replicates.

Mj-NEROSs protein has a predicted architecture comprising several domains (**Fig. 2b**). Following a signal peptide (SP) for secretion from the nematode to the plant, a cysteine-rich, proline-rich domain containing a chloroplast transit peptide (cTP) was identified. At the C-terminus of Mj-NEROSs a ground-like domain was found. The ground-like domains have been reported to modulate the activity of membrane proteins in *Caenorhabditis elegans* ^50^. The tomato ISP, on the other hand, contains a chloroplast transit peptide (cTP), Fe-S domain, and a Rieske domain (**Fig. 2b**). To gain insight into the structural basis of their interaction, we created deletion constructs of both genes. The first fragment of ISP consisted of the Fe-S and cTP domain and the second of the Rieske domain. We found that NEROSs protein binds and recognizes particularly the Rieske part of ISP (**Fig. 2c**), but the interaction is stronger with the full-size ISP (**Supplementary Fig. 2b**). On the other side, despite splitting NEROSs into several parts, all parts together with the full-length ISP grew on the selective medium (-LWH), thus showing that they all contribute to the more robust interaction with the full-length ISP protein (**Supplementary Fig. 2c**). In addition, pairwise Y2H suggests that NEROSs might not act as a monomer but rather a dimer or a multimer. The dimeric and possibly multimeric interactions are established via N-N terminus binding or C-C terminus interaction, but not between N and C terminal parts (**Supplementary Fig. 2d**). Taken together, we discovered that the *M. javanica* effector NEROSs acts as a homodimer and interacts with the Rieske part of the tomato chloroplast protein ISP.

### *In planta* validation of NEROSs-ISP interaction in the chloroplast

To validate the interaction between ISP and NEROSs we used bimolecular fluorescence complementation (BiFC). NEROSs and ISP were fused to N and C terminal parts of mVenus fluorescent protein, respectively. Their co-expression reconstituted mVenus fluorescence in the chloroplasts of agro-infiltrated *N. benthamiana* leaf cells (**Fig. 3a**). In contrast, there were no fluorescent signals when the chloroplast localization signal was fused to C and N terminal parts of mVenus and tested against NEROSs-NmVenus or ISP-CmVenus, respectively. The observed fluorescence indicates that NEROSs and ISP interact *in planta*. Moreover, by using a chloroplast Rubisco marker we showed that this interaction happens explicitly in the chloroplast (**Fig. 3a**), which was not surprising considering that both proteins possess a chloroplast targeting signal. However, we wondered if the formation of the NEROSs-ISP complex is a requirement for chloroplast localization. To this end, C-terminal fluorescent fusions to both proteins were created and their localization analyzed individually. Both NEROSs-eGFP and ISP-mRFP, when infiltrated separately (**Fig. 3b**) or together (**Supplementary Fig. 3a**), localized in the chloroplast. This localization is observed with different fluorescent tags, excluding the possibility of labeling artifacts (**Supplementary Fig. 3b**). We hypothesize that this interaction could happen only if both proteins have already been transported to chloroplasts. To test this hypothesis, we introduced mutations within NEROSs’ cTP. Using the LOCALIZER tool, the possible substitutions that would lead to the loss of chloroplast localization were narrowed down to 3 amino acids. Indeed, chloroplast localization was lost with NEROSs_S51D_L52D_F54D_, here named NEROSs_cTPmut_ (**Fig. 3c**). Notably, this mutant also lost its ability to interact with ISP *in planta*, as shown in a BiFC assay (**Fig. 3d**). Putting the fluorescent tag on the N-terminal part of NEROSs had the same effect (**Supplementary Fig. 3c**), probably because this masks cTP thus preventing the transport of NEROSs to the plant organelle. These results are further strengthened since when at least one of the proteins is tagged at its N-terminus, this did not yield the reconstruction of a full mVenus signal in our BiFC assays (**Supplementary Fig. 3d**). We conclude that the interaction between ISP and NEROSs *in planta* happens when both proteins reside in the chloroplast.

**Fig. 3.**
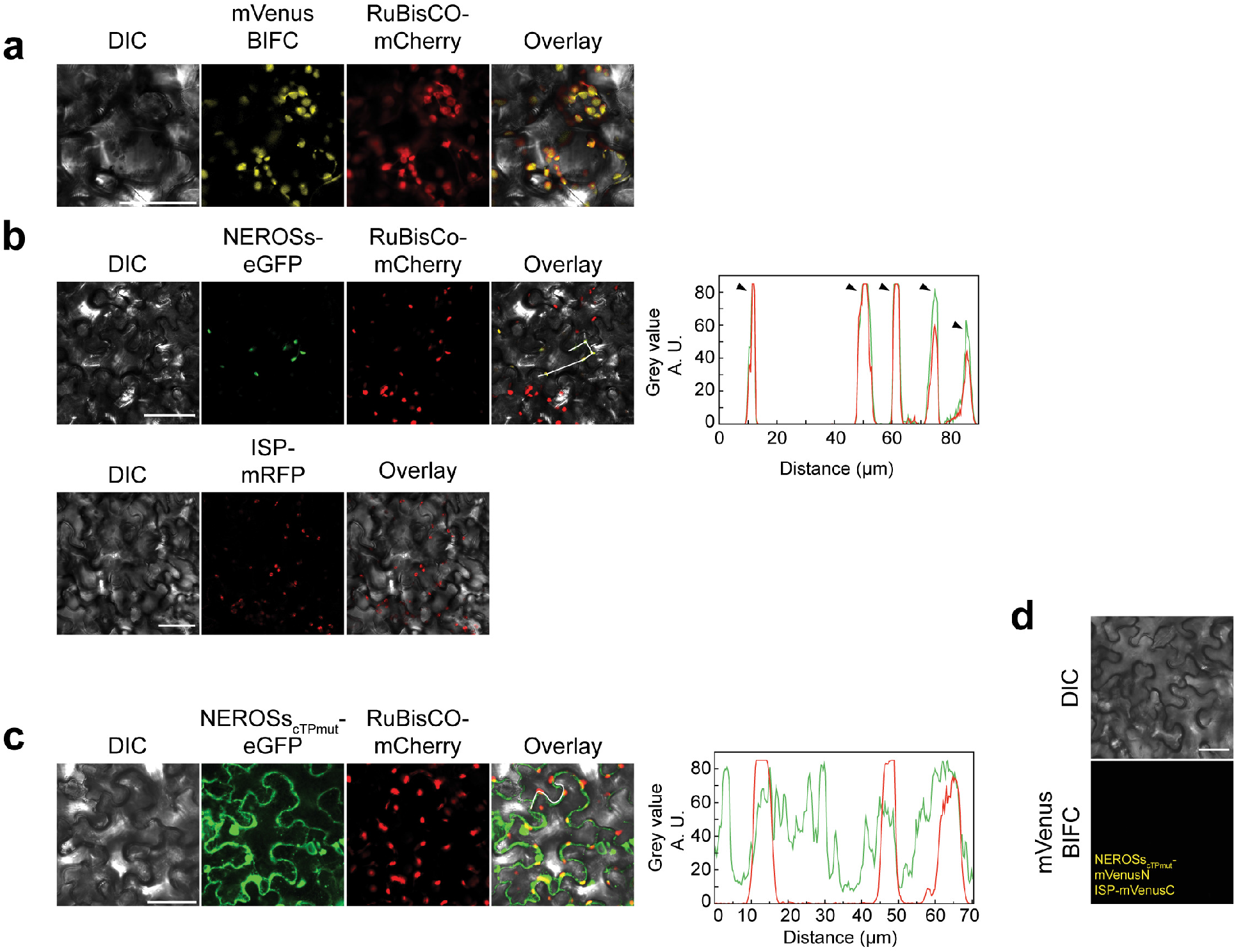
NEROSs and Sl-ISP localize and interact in chloroplasts. **a** Detection of the NEROSs-ISP interaction in chloroplasts by bimolecular fluorescence complementation (BIFC) assays in *N. benthamiana* leaves transiently expressing the marked constructs; RuBisCO-mcherry is used as a chloroplast marker. **b** NEROSs and ISP show chloroplast localization. Top: representative differential interference (DIC) contrast and fluorescence images of leaf tissues of *N. benthamiana* transiently expressing NEROSs-eGFP or RuBisCO-mcherry; fluorescence intensity profiles of the corresponding signals in the same cells; arrows indicate their colocalization. Bottom: same for ISP-mRFP. **c** NEROSs mutant does not localize in the chloroplasts. From left to right: representative DIC contrast and fluorescence images of leaf tissues of *N. benthamiana* transiently expressing NEROS_scTPmut_-eGFP or RuBisCO-mcherry; fluorescence intensity profiles of the corresponding signals in the same cells. **d** NEROSs mutant does not interact with ISP. No signal is detected by bimolecular fluorescence complementation (BIFC) assays of NEROS_scTPmut_ and ISP in *N. benthamiana* leaves transiently expressing the marked constructs; fluorescence intensity profiles of the corresponding signals in the same cells. Scale bars 50 μm.

### NEROSs hinders the plant basal immune response by suppressing ROS, callose deposition and HR

Next, the physiological effect of the ISP-NEROSs interaction in plants was investigated. Because the interaction of the two proteins is confirmed to happen in chloroplasts, where ISP, part of the cytochrome b6f-complex, plays a vital role in a photosynthetic relay connecting PSII to PSI ^51^, the interaction with NEROSs might interfere with the electron transport ^52^. To test this hypothesis, the effect of transient expression of NEROS*s* on the electron transport rate (ETR) in tobacco was measured. As shown in **Fig. 4a**, overexpression of *NEROSs* in infiltrated *N. benthamiana* leaves led to a significant decrease in ETR at 72 HPI (hours post infiltration) compared to the eGFP control.

**Fig. 4.**
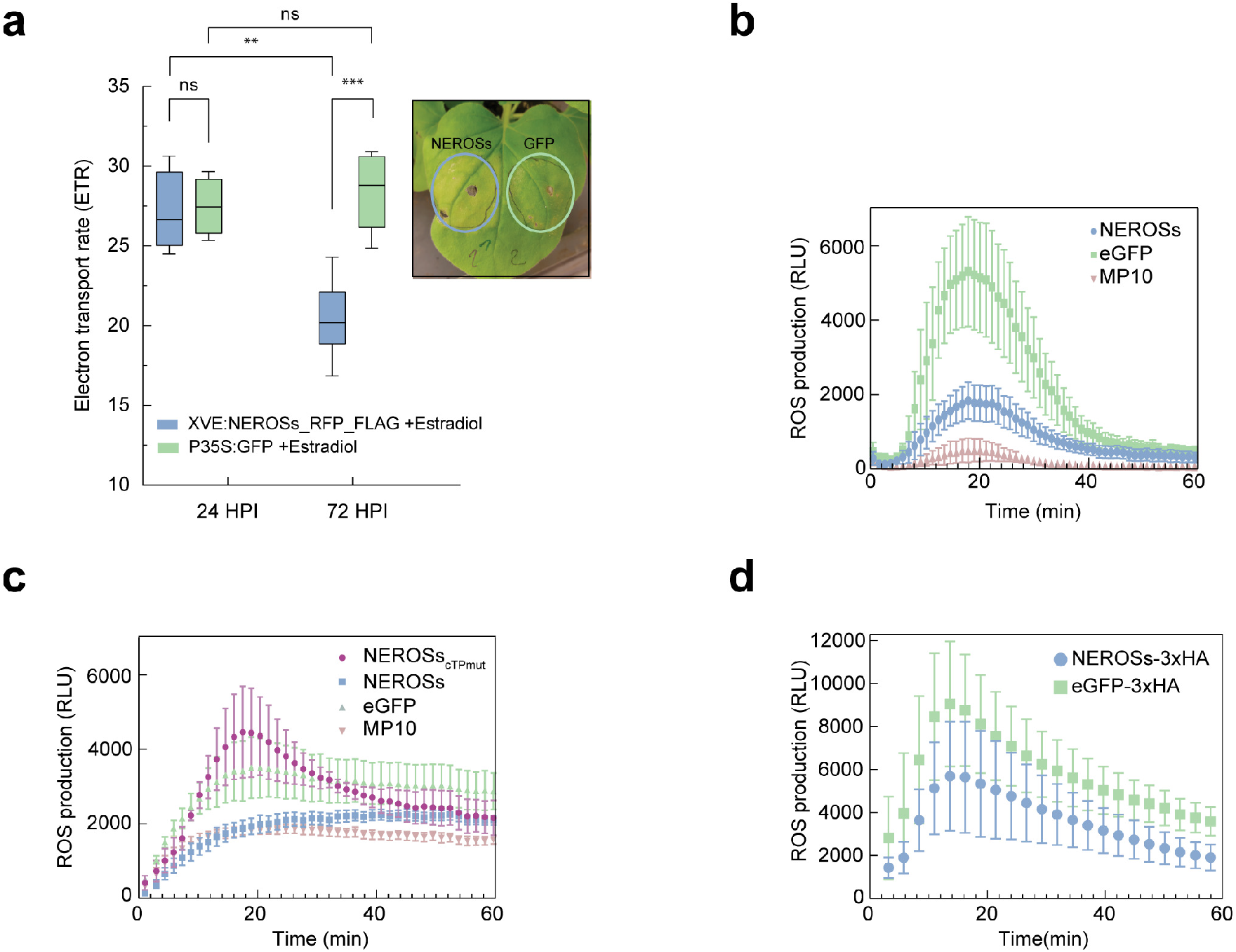
NEROSs suppresses ETR and ROS production in leaves and ROS in roots. **a** NEROSs suppresses the electron transport rate (ETR) in *N. benthamiana* leaves infiltrated with Agrobacterium expressing NEROSs-RFP-Flag or eGFP (as the control) from the estradiol inducible promoter; the measurements were performed at two time points: 24h and 72h post infiltration (HPI); the mean and standard deviation were calculated from six independent replicates; the asterisk indicates a significant difference (ns non-significant,** P < 0.01, *** P < 0.001 unpaired two-tailed Student’s t-test with Welch’s correction (at 72 HPI, t=5,808, df=9,972)). **b** ROS accumulation in chloroplasts was reduced in leaves transiently expressing NEROSs. Tobacco leaves infiltrated with *Agrobacterium* expressing NEROSs, eGFP (as the negative control) or MP10 (as the positive control) were tested for ROS production. **c** Same as in *b* with NEROSs mutant (purple) that does not localize in the chloroplast and does not interact with SI-ISP. **d** ROS production in the tomato hairy roots was reduced when NEROSs is transiently expressed. The tomato hairy roots expressing NEROSs-3xHA, eGFP-3xHA (as the control) were tested for ROS production. The number of leaf discs analyzed in all ROS measurements was 24. For roots, 4 independent biological replicates and 8 technical were used. For *b, c and d*, bars indicate the mean and standard deviation; a two-tailed Student’s t-test was used to determine significant differences per time.

Under unfavorable conditions, electron leakage from PSI to the Ferredoxin complex leads to the accumulation of O_2_^-^, which undergoes dismutation to H_2_O_2_^53^. To assess the H_2_O_2_ rate in the presence of NEROSs, ROS production was induced by flg22 in *N. benthamiana* leaves expressing Mp10, eGFP or NEROSs. Mp10 was included as a positive control as it was previously shown to suppress the oxidative burst induced by flg22 ^54^. We found that NEROSs suppresses the flg22-induced ROS production at 72 HPI in *N. benthamiana* leaves, while this was not the case with the negative control eGFP (**Fig. 4b**). As described above, mutations in the chloroplast transit peptide from NEROSs prevented its chloroplast localization and subsequently its interaction with ISP. To test whether the decrease in ROS accumulation is a consequence of the specific NEROSs localization and thus protein-protein interaction in the chloroplast, the ROS assay was repeated using NEROSs_cTPmut_. This mutant failed to suppress ROS production similarly to the eGFP (**Fig. 4c**), strengthening the proposed action where NEROSs is responsible for hijacking the ISP function away from its plant defense role in chloroplasts.

A callose deposition assay was performed as one of the landmarks of PTI responses to confirm that interference with ROS production is part of PTI suppression. Upon expression of NEROSs in the *N. benthamiana* leaves and treatment with flg22, we observed less callose deposition in the NEROSs sample compared to GFP or NEROSs_cTPmut_ (**Supplementary Fig. 4a**), suggesting once again that specific localization of NEROSs is important for suppression of PTI.

The RKN effector, NEROSs, should play its role at the site of the infection, which are the tomato roots. Previously, it has been reported that chloroplast-like structures can be found in giant cells ^2^, and the expression of genes involved in chloroplast biosynthesis increases upon RKN infection ^3^. In addition, the existence of ISP was shown not just in chloroplasts but also in non-photosynthetic plastids ^55^. Keeping in mind the conservancy and the essentiality of the cytochrome b6f complex ^56^, we suspect that similar processes and systems exist in root cells with chloroplast-like structures. If this is true, inducible NEROSs-3xHA expression in hairy roots could suppress ROS production. In fact, the expression of NEROSs-3xHA led to the decrease of ROS in hairy roots after flagellin treatment compared to the eGFP-3xHA (**Fig. 4d and Supplementary Fig. 4b**). Finally, NEROSs can also suppress cell death triggered by INF1 (**Supplementary Fig. 4c**).

### Transcriptional changes revealed by RNA-seq

To broaden our understanding of NEROSs functioning in plant cells, the effect of NEROSs on tomato gene expression was investigated in hairy roots. RNA was extracted from hairy root lines 24 hours upon the induction of NEROSs-3xHA, NEROSs_cTPmut_-3xHA, or eGFP-3xHA (**Supplementary Fig. 4b**). Then, RNA was sequenced using Illumina high throughput sequencing technology. Analysis of differentially expressed genes (DEG) identified 308 up- and 402 down-regulated genes for NEROSs compared to eGFP control lines, or 934 up- and 558 down-regulated genes for NEROSs compared to NEROSs_cTPmut_ **(Supplementary Table 3 and 4)**. qRT-PCR on seven randomly selected (but with a similar trend in both comparisons) up-regulated and down-regulated genes showed an appropriate correlation with the transcriptome data (**Supplementary Fig. 5a, 5b**, and **Supplementary Table 5**). When NEROSs was compared to the eGFP control, gene ontology (GO) term analysis highlighted an over-representation of downregulated genes involved in “protein folding or unfolding under stress conditions”, as well as “response to hydrogen peroxide and ROS” (**Fig. 5a** and **Supplementary Table 6**). The latter fits well with all our previous observations. Similar results were obtained when NEROSs was compared to the mutant that lost its chloroplast localization and its interaction with ISP *in planta*, further supporting that the NEROSs plastid localization and possibly its interaction with ISP represent the primary mode of its action. In addition, we observed defense responses being downregulated (**Fig. 5b** and **Supplementary Table 8**). Among the genes uniquely downregulated in NEROSs compared to the mutant (**Supplementary Fig. 5c and Supplementary Table 11**) but not compared to eGFP we identified chloroplast-related genes as well as those involved in photosynthesis (**Supplementary Fig. 5d**). Interestingly, comparing NEROSs expressing hairy roots with NEROSs_cTPmut_ revealed: “hydrogen peroxidase catabolic processes”, “oxidoreductase processes,” as well as “cell wall-related genes” among those being up-regulated (**Fig. 5c, d** and **Supplementary Table 7, 9**). On the other hand, we did not find significant or new GO terms among other unique genes (**Supplementary Fig. 5c**).

**Fig. 5.**
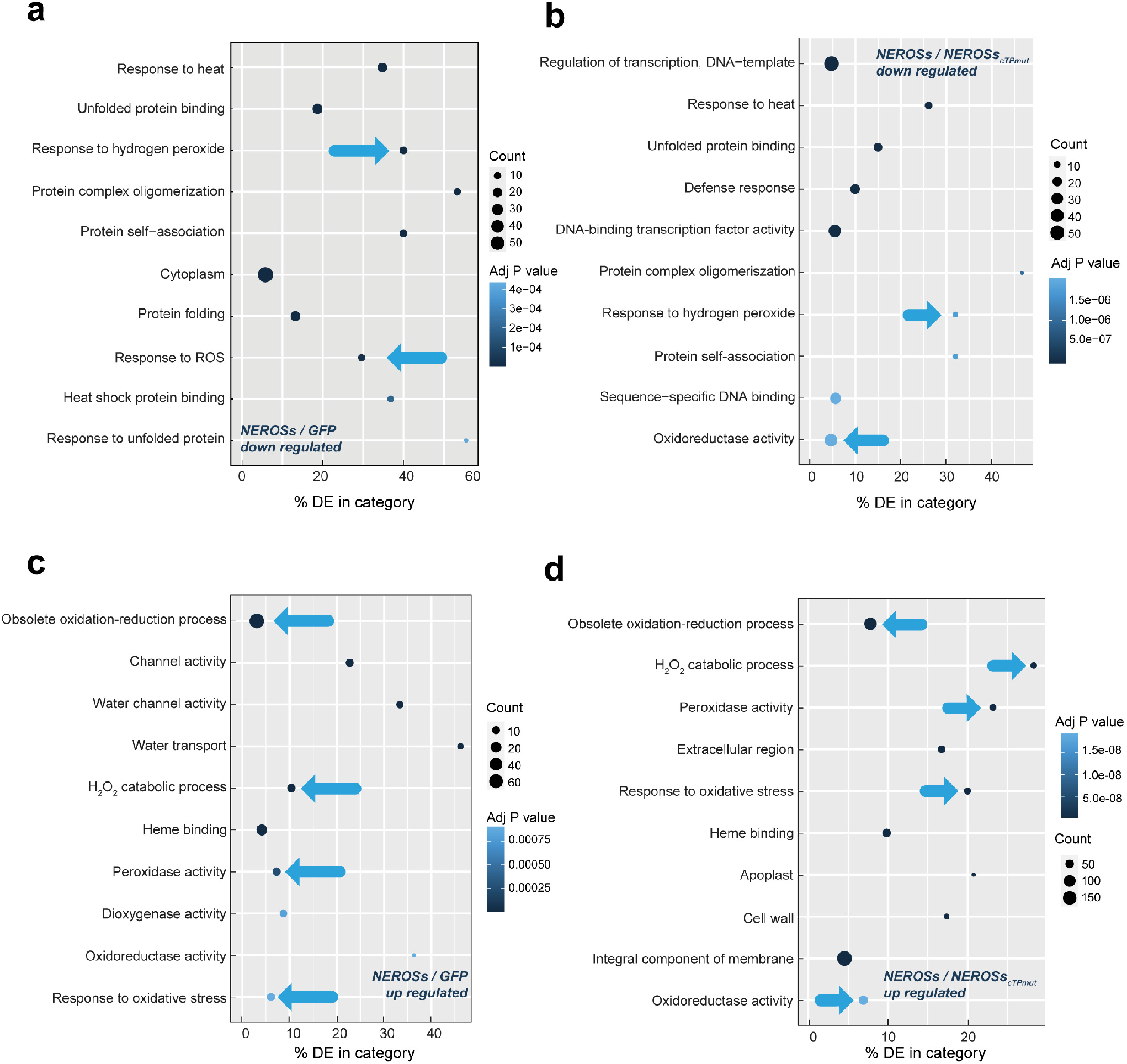
Transcriptional changes caused by NEROSs in tomato hairy roots. **a** Identification of differentially expressed genes (DEGs) by GO enrichment in the tomato hairy roots expressing NEROSs after transformation by *Agrobacterium*. Bubble Plot of top ten overrepresented gene categories downregulated in NEROSs compared to eGFP (the control). **b** same as *a* for NEROSs compared to NEROSs_cTP_ (the mutant). **c** Bubble Plot of top ten overrepresented gene categories upregulated in NEROSs compared to eGFP. **d** same as *c* for NEROSs compared to NEROSs_cTP_. GO-seq tool was used for analysis; blue arrows indicate processes involved in ROS pathway.

## Discussion

Many different biotrophic pathogens challenge plants. Nevertheless, they share some common features, such as producing effector proteins to overcome plant defense mechanisms and take the lead in the ongoing evolutionary battle ^57,58^. Therefore, to comprehend and find strategies to boost plant immunity and improve yield, it is essential to understand the mechanism of the effectors’ mode of action and their targets.

In this work, we focus our research on the root-knot nematode (RKN) effector, Mj-NEROSs, and demonstrate its versatile role, finally leading to the suppression of the immune system. While previously shown that silencing the nearly identical *M. incognita* effector has a significant impact on nematode life cycle completion ^42^, the molecular mechanism behind it remained unknown. Our work reveals that Mj-NEROSs is an effector protein expressed highly during mid-later parasitic stages (**Fig. 1a**), contrasting with the previously reported dominant expression in the egg stage (Joshi *et al*., 2020). By analyzing more intermediate stages, broader insight was obtained into its temporal expression. Indeed, expression in the eggs is higher than in the nematodes at 30 DPI, but the expression peaks at 18 and 21 DPI. Most of the reported RKN effectors are produced in dorsal or subventral glands ^5^. Herein, combining ISH and immunohistochemistry, we provide insight into the spatial and temporal localization of *Mj-NEROSs* transcripts and protein, confirming that this effector is produced in the subventral esophageal glands (**Fig. 1b, 1c**). Moreover, we found that the nematode secretes NEROSs explicitly in the giant cells, concentrated in giant cell foci (**Fig. 1c** and **Supplementary Fig. 1b**). While other nematode effectors have been localized, most of them target the nuclei, cytosol, or the apoplast ^13,59,60^. Hence, this is the first study where such localization of giant cell foci has been reported. We speculate that these foci present plastid organelles in giant cells. Because of the lack of artificial dyes available for plastid organelles and the inability to transform the nematodes, molecular genetic approaches were combined with confocal microscopy to study the localization of this protein further after the expression of the effector *in planta*. We showed that giant cell foci are most likely plastids and that the presence of an intact chloroplast targeting signal in Mj-NEROSs is necessary for its localization in these organelles (**Fig. 3b, 3c, and Supplementary Fig. 3b**). Notably, a chloroplast localization signal was found in many other RKN effector proteins (**Supplementary Table 1**), attesting to the importance of these organelles in giant cells for RKN-plant interaction and raising the question of what is the role of all these effectors in this single compartment. This question remains to be tackled in future studies.

In alignment with these analyses, NEROSs was found to suppress ROS production and to interfere with plant defense only when present in plastids, either in roots or shoots (**Fig. 4c, 5b and Supplementary Fig. 4a**). The ISP protein, which also localizes in chloroplasts (**Fig. 3b**), was identified as a direct partner (**Fig. 2a, 2c, 3a** and **Supplementary Fig. 2a, 2b**). Not only NEROSs interacts with ISP, but it can also suppress the electron transfer rate, ETR (**Fig. 4a**), the primary function that ISP plays in the plastids (**Fig. 6** and ^51^), thus indicating a possible link between its direct target and its role in ROS suppression.

**Fig. 6.**
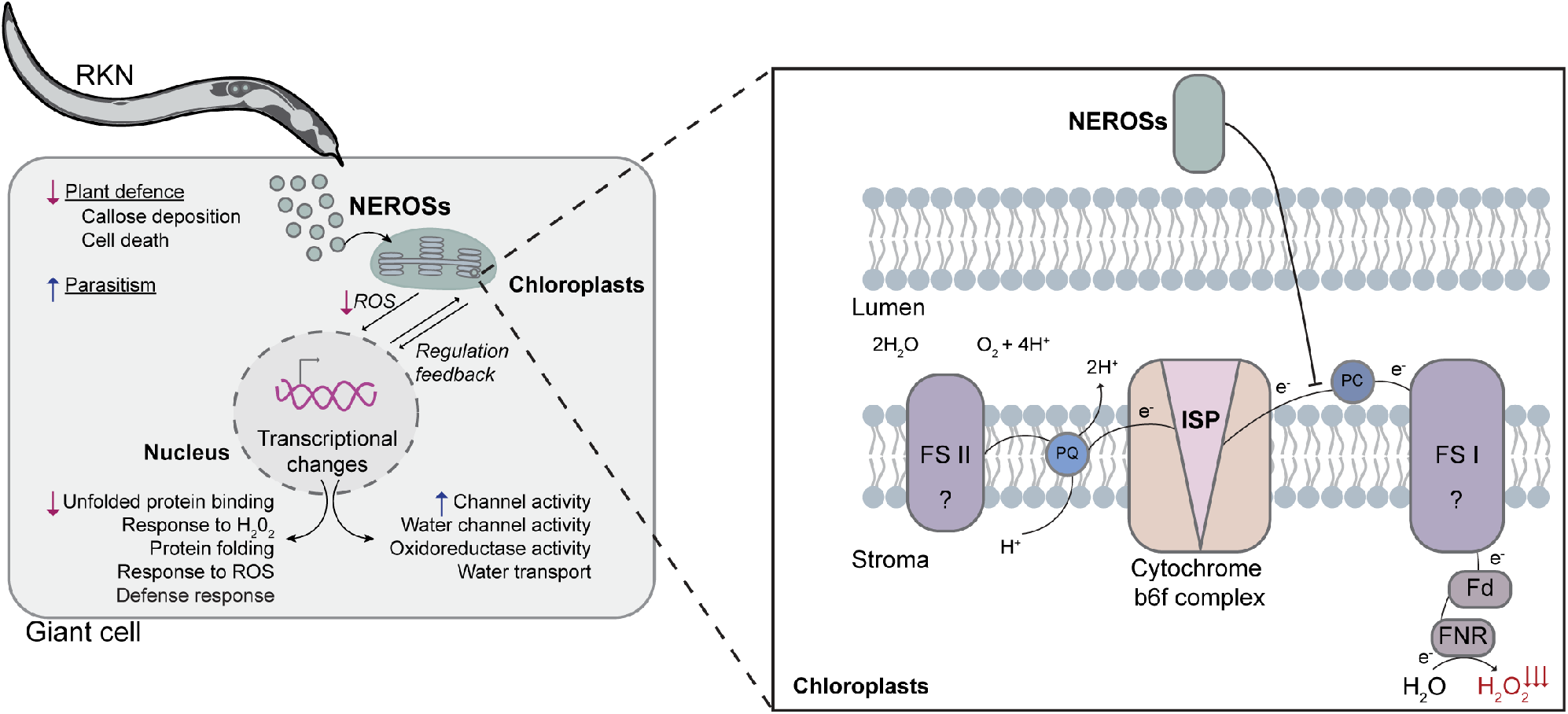
Model for the function of Mj-NEROSs during plant-nematode interaction. RKNs secrete the effector NEROSs from their sub-ventral esophageal gland into the giant cells where NEROSs interacts with Sl-ISP in the plastids. The formed complex interferes with the function of this Sl-ISP in the electron transport chain and results in a decrease of the electron transport rate, which causes the attenuation of ROS accumulation in these organelles (zoom). Lower plastid ROS production influences redox communication to the nucleus and thus influences gene expression. Primarily, genes involved in response to ROS, defense response, and protein folding are downregulated, but there is also an upregulation in the expression of genes involved in channel activity and water transport activity. Overall, changes caused by NEROSs lead to reduced plant defense and most likely increase nematode parasitism.

Previously during RKN infection of tomato plants, it has been observed that the *ISP* gene has been upregulated at the early stages, while at later stages, the expression goes down ^61^. Interestingly, the time-point of the *ISP* expression switch roughly overlaps with Mj-NEROSs/*Mj-NEROSs* peak of expression (**Fig. 1a, 1c**, and **Supplementary Fig. 1a**). Consistent with the hypothesis that NEROSs could be directly involved in regulating *ISP* expression, we observed downregulation of *ISP* mRNA in roots expressing NEROSs (**Supplementary Table 3**), most likely as a possible feedback mechanism for the downregulation of ROS from plastids (**Fig. 6**). Our results, hence, point out that Mj-NEROSs suppresses the function of ISP on the protein level by protein-protein interaction (**Fig. 2a**) but also indirectly by lowering the transcription of the *ISP* gene.

It is a common feature of many pathogenic effectors to inhibit cell death triggered by plant defense mechanisms ^17,62^. NEROSs has the same function, pointing further to its direct role in plant immunity (**Supplementary Fig. 4c**). Keeping up with the theme, we can now claim that the biological effect of NEROSs is threefold: at the protein level, it interacts with ISP in plastids; at the physiological level, it interferes with the electron transport rate and hence ROS production; at the transcriptome level, it can alter the expression of the genes involved in those processes.

To explain our discovery that NEROSs interacting with ISP blocks its role in the electron transport chain, as well as the ROS production in chloroplasts and plastid-like structures in giant cells, we propose that a decrease in ROS production could lead to the overall suppression of plant immunity, where the plant stays defenseless on different fronts against RKN. It cannot be excluded that Mj-NEROSs has multiple binding partners that were not identified in our screen. The latter could explain the complex transcriptome and physiology changes observed in our RNA-seq data. The development of protocols for nematode transformation to tightly follow the dynamic of the effector in combination with multiple proteomic techniques to screen for interactions will likely shed light on the unresolved mystery of complex transcriptional changes. Unraveling the spatiotemporal expression and molecular mode of action of the effector, the importance that root plastids play in plant defense was touched upon for the first time. With this, an arsenal of questions has been opened, such as why and how they can be transformed into defense organelles and testify their importance as effector’s targets. But also, besides ISP, what are the other players in the electron transport chain and ROS production in root plastids?

Answering these questions will impact either designing new chemicals that target those interactions for further yield improvement or purely fundamental on understanding organelle changes in the complex interkingdom interaction and assigning them novel functions. Altogether (**Fig. 6**), our data illustrate that NEROSs produced in the subventral esophageal glands is being secreted to giant cells and going to the plastids. In plastids, it targets ISP blocking its ability to transfer electrons and further inhibiting ROS production. NEROSs lowers ROS production in the plastids, which leads to overall reduced ROS levels in the cell. Thus, it alters the expression of redox-related genes and interferes with the recognition of misfolded proteins. Overall, this leads to decreased plant defense, which is reflected in reduced hypersensitive response and callose deposition and enhances pathogen survival.

## Materials and Methods

### Strains

*Meloidogyne javanica* and *Meloidogyne incognita* morelos cultures were maintained using a susceptible tomato (*Solanum lycopersicum* cv Moneymaker) in two-liter plastic pots containing potting soil (Structural, Kaprijke, Belgium). Infected tomato plants were grown in a greenhouse under 8/16 h night/day conditions at 22-24 °C and 16 °C, respectively. *Nicotiana benthamiana* plants were grown in a greenhouse under 8/16h night/day conditions at 28 °C and 16 °C, respectively. *Escherichia coli* (Top10) and *Agrobacterium tumefaciens* GV3101 were routinely grown in LB/YEB medium at 37 °C and 28 °C, respectively. The yeast strain *Saccharomyces cerevisiae* PJ69-4α was cultured at 30 °C for Y2H assays as described below. All the plant extracts, tissues, and bacterial and yeast strains were stored at −80 °C.

### *In silico* sequence analysis

Using Meloidogyne INRAE Blast (https://meloidogyne.inrae.fr/), we found the closest homolog of Mi-4D01 (Accession number AF531162) ^5^ to be Mjav1s02606g024064 (M.Javanica Scaff2606g024064), here referred as Mj-NEROSs. SignalP 5.0 was used to predict signal peptides. For all the experiments, LOCALIZER ^63^ was used to predict the protein localization and chloroplast localization signal. Motif scan and NCBI Conserved Domain Search were used to predict motifs and domains in the proteins.

### Plasmid and strain construction

All plasmids used in this study are listed in **Supplementary Table 2** and constructed as indicated in **Supplementary Table 2**. Standard molecular cloning methods were used, and DNA assembly was performed using Gateway cloning, Gibson assembly or site directed mutagenesis. All oligos used in this study are listed in **Supplementary Table 2**.

Tomato (Solyc12g005630), tobacco (Niben101Scf21348g00006), and Arabidopsis (AT4G03280) ISP coding sequences were used as templates for amplification from cDNAs. **Stage expression analysis by qRT-PCR**. To analyze the transcripts of Mj-NEROSs during *M. javanica* development, RNA samples were prepared from 200 *M. javanica* nematodes at different life stages, as indicated with the RNA prepmicro kit (Tiangen). The cDNA was then synthesized using the TransScript One-Step gDNA Removal and the cDNA Synthesis SuperMix kits (Transgen Biotech). qRT-PCR was performed using the THUNDERBIRD SYBR® qPCR Mix (TOYOBO) and operated in a Takara TP800 real-time PCR system. Two primers q4D01-F/q4D01-R and qMjact-F/qMjact-R were designed to amplify the Mj-NEROSs and internal control gene Mj-β-actin (accession no. AF532605). The relative changes in gene expression were determined using the 2^-ΔΔCT^ method. Three independent biological replicates were performed, each biological replicate comprising three technical replicates.

### ISH (*in-situ* hybridization)

The experiment was performed as previously described^64^. Briefly, a segment of *Mj-NEROSs* was amplified (primers 4D01-ish-F and 4D01-ish-R) and used as a gene-specific probe. A digoxigenin (DIG)-labeled antisense (AS) probe was synthesized using the T7 polymerase (PROMEGA), for hybridization on pre-parasitic stage 2 juvenile nematodes (ppJ2). The results were examined using a Nikon ECLIPSE Ni microscope (Nikon, Tokyo, Japan). The experiment was repeated three times, illustrating the same *Mj-NEROSs* AS probe localization.

### Peptide synthesis and antibody purification

Peptides used for immunolocalization of Mj-NEROSs were designed according to the Thermo Fisher Scientific Antigen profiler. The designed peptides were blasted against the tomato genome SL3.0 using Solgenomics Basic Local Alignment Search Tool (https://solgenomics.net/tools/blast/.) To avoid the background, we checked for potential similarities between the selected peptide sequences and host plant proteins. Additionally, WormBase ParaSite Blast (https://parasite.wormbase.org/Multi/Tools/Blast) and Meloidogyne INRAE Blast (https://meloidogyne.inrae.fr/) were used to check for potential background among nematode proteins. The last selection of immunogenic peptides was made based on the Alfa2fold predicted 3D protein structure ^65^. Two rabbits were immunized by injecting two synthetic peptides: GRRKRESGA and STKSGIKRIGEEKN (Thermo Fisher Scientific). Sera before peptide co-injection and 72 days after co-injection were used as primary antibodies to check for specific bands in infected compared to non-infected roots. Primary antibodies were purified from the rabbit with the lowest background (Thermo Fisher Scientific).

### Gall fixation and immunolocalization

Four weeks-old tomato plants grown in pots were infected with freshly hatched ppJ2s. Galls were then dissected from infected root systems at 7-, 14-, 21-, and 40 days post infection (DPI), washed thoroughly with water and placed in fixative (4% formaldehyde in 50mM PIPES buffer, pH 6.9). Galls were then vacuum treated for 30 min to remove air from the tissue, transferred to fresh fixative in cell strainers, and placed at 4 ºC, shaking gently for 1-3 weeks (refreshing weekly), depending on the gall size. Fixation and embedding were further performed as previously described ^66,67^. Galls were then sectioned to 5 μm using an microtome and placed on poly-lysine-coated slides ^68^. The immunolocalization was performed according to ^66,68^ slightly modified. Gall sections were incubated with a blocking solution (BS, 1% bovine serum albumin (BSA) in 50mM PIPES buffer (pH 6.9) at RT for 30 min. The Mj-NEROSs-specific antibody was diluted to 1:50 in fresh BS, and 250 μl was placed on each slide. BS, as well as dilutions containing the antibody were centrifuged for 5 min and incubated for 1 h at 37 ºC. As negative control, 250 μl fresh blocking solution were added to each slide and only treated with secondary antibodies. All slides were incubated in a humid box at 4 ºC overnight and then placed at 37 ºC for 1 h. Slides were then washed for 30 min in PIPES buffer and subsequently incubated for 2 h at 37 ºC with the secondary antibody (goat anti-rabbit IgG (H+L) Alexa Fluor 488, Thermo Fisher Scientific), diluted 1:300 in BS. Finally, slides were washed in PIPES buffer for 30 min and nuclei were stained with 1 μg/ mL 4,6-diamidino-2-phenylindole (DAPI) for 5 min at RT, rinsed in distilled water, mounted in 90% glycerol, and cover slipped. Sections were observed with a fluorescence microscope (Zeiss, Oberkochen, Germany) and using a double bandpass filter (Zeiss) auto-fluorescence was visualized in red and Alexa-488 fluorescence in green. In addition, differential interference contrast (DIC) transmission and DAPI-stained DNA images were obtained to provide overlaid images.

### Yeast-two hybrid

Y2H analysis was performed as described in ^69,70^, with the GAL4 system. Briefly, bait and prey open reading frames were fused to the GAL4-AD or GAL4-BD via cloning into pGADT7 or pGBKT7, respectively. The *S. cerevisiae* PJ69-4α yeast strain was co-transformed with bait and prey using the polyethylene glycol (PEG)/lithium acetate method. Transformants were selected on Synthetic Defined (SD) media lacking Leucine and Tryptophane (Clontech). Three individual clones were grown ON in liquid cultures (lacking Leucine and Tryptophane) at 30°C and 10x or 100x dilutions were spotted on the control media (SD-Leu-Trp) and selective media lacking Leu, Trp, and His (Clontech) with and without the addition of 3-Amino-1H-1,2,4-triazole (Acros Organics (Thermo Fisher Scientific)) to test the strength of the interaction.

### BiFC and cell imaging

For microscopy, all constructs (**Supplementary Table 2**) were transformed into *A. tumefaciens* strain GV3101 and complementary constructs were co-infiltrated into *N. benthamiana* leaves at OD600 final concentration of 0.3 as described above. Infiltrated plants were kept in daylight at RT and, 48 h after inoculation, the infiltrated spots were imaged using a confocal laser-scanning microscope (Nikon Instruments Inc., Tokyo, Japan). eGFP was excited with a wavelength of 488 nm and emission was detected at 495– 530 nm; eRFP was excited with a wavelength of 561 nm and emission was detected at 592– 632 nm. Auto-fluorescence of chlorophyll was collected at 657–737 nm.

For BiFC, ISP was fused with the C-terminal fragment of mVenus ^71^ and NEROSs to the N-terminal fragment of mVenus. Vectors with only the N or C-terminal fragment of mVenus ^71^ were used as a control as well as the cTP (chloroplast localization signal) fused to C or N-terminal fragment of mVenus (cTP-mVenusC or cTP-mVenusN). Constructs were transformed into *A. tumefaciens* strain GV3101 and complementary constructs were co-infiltrated into *N. benthamiana* leaves at OD600 final concentration of 0.3. The mVenus signal was analyzed 72 h after infiltration. mVenus was excited with a wavelength of 514 nm and emission was collected at 530–575 nm using confocal microscopy (Nikon Instruments Inc.)

### Electron Transport Rate

Electron transport rates (ETR) were derived from chlorophyll fluorescence measurements performed with a portable photosynthesis system (LI-6800, Li-COR, Lincoln, Nebraska USA) according to the manufacturer’s instructions (https://www.licor.com/env/support/LI-6800/topics/fluorometer-experiments.html, ‘Determine PSII efficiency’). NEROSs-RFP-Flag and GFP were transiently expressed in 4-week-old tobacco (*N. benthamiana*) leaves and the expression was induced using 100 μM β-Estradiol (cat. no: E8875-1G; Sigma). Leaves were dark-adapted for 20 min before chlorophyll fluorescence and ETR were measured at 20°C. Measurements were done on 6-8 biological replicates.

### ROS production assay in *N. benthamiana* leaves

The *Agrobacterium* carrying *NEROSs* or *NEROSs*_*cTPmut*_ under control of the 35S promoter (vector pK7WG2 **Supplementary Table 2**) was grown for 2 days at 28ºC in LB medium with 25 μg/ml rifampicin, 25 μg/ml gentamicin, or 100 μg/ml spectinomycin. A plasmid expressing eGFP was used as a negative control; the aphid (*Myzus persicae*) effector Mp10 was used as a positive control ^72^ Flg22 (cat. no. RP19986, GenScript) was used as an inducer of ROS production. The protocol was performed as described before ^13^. Briefly, before infiltration, the overnight *Agrobacterium* cultures were washed and diluted in infiltration buffer (10mM MgCl_2_, 10 mM MES and 0.2mM acetosyringone), to reach OD_600_ of 0.3 and incubated for three hours in the dark at room temperature. The next day, *N. benthamiana* 5-6 weeks old leaves were infiltrated with the Agrobacteria using a needleless syringe. The appropriate Agrobacteria were infiltrated at different spots on the same leaf. For each assay, one leaf from eight plants was used with two replicates per spot. Approximately 30 h after infiltration, 16 mm^2^ leaf discs were collected from the infiltrated areas with a cork borer, transferred to 96-well plates, and floated overnight on 200 μl filter-sterilized ultrapure water for recovery. After 48 h, the water was removed and replaced by a mixture of 100 nm flg22 (QRLSSGLRINSAKDDAAGLAIS^73^), 0.5 mM luminol probe 8-amino-5-chloro-7-phenyl-pyrido[3,4-d] pyridazine-1,4(2H,3H) dione (L-012) (Wako Chemicals, Richmond, Virginia, USA) and 20 μg/ml horseradish peroxidase (Sigma, Saint Louis, Missouri, USA). ROS production was measured by a luminol-based assay ^74^ over 60 min with integration at 750 ms.

### ROS production assay in hairy roots

To measure ROS production in hairy roots, we adopted and modified the protocol ^75^. Briefly, the transformed tomato hairy roots were used (pKCTAP vector, **Supplementary Table 2**). Approximately 1 cm (+/-0.05cm) roots were inoculated in Murashige and Skoog (MS) liquid media with 100 μM estradiol for 24 h in 96-well plate. MS liquid media were removed and 100 μM estradiol in ddH2O was added for 12 h. Afterward, water was removed and a 200 μl solution containing 3μg/ml horseradish peroxidase, 0.5mM luminol 8-Amino-5-chloro-2,3-dihydro-7-phenyl-pyrido [3,4-d] pyridazine sodium salt (L-012, Wako Chemicals) and 50nM flg22 was added.

Measurements were carried out in white nontransparent 96-well plates for 60 min. Each treatment contained 4 biological and 3 technical replicates. ROS production was measured by a luminol-based assay ^74^ over 60 min with measurements every 160 s with integration at 750 ms.

### Cell death assay

*A. tumefaciens* strain GV3101 was used for all constructs (see **Supplementary Table 2**); plasmid pK7WG2-*GFP* was used as a negative control.

Agrobacteria carrying a plasmid for expression of *NEROSs* or *INF1* ^76^ were grown for 2 days in 10 ml of LB medium with the appropriate antibiotics. Before infiltration, the cells were pelleted, washed, resuspended in infiltration buffer (10mM MgCl_2_, 10 mM MES and 0.2mM acetosyringone), and incubated for at least 3 h at RT. Depending on the combination of constructs, the final concentration in the mixtures was adjusted to an OD_600_ of 0.5. The mixtures were spot infiltrated into *N. benthamiana* leaves of 5–6-week-old plants. The negative controls were infiltrated on the same leaf as the tested effector. For each plant, two leaves were infiltrated, and 20 plants were used per assay. The response was recorded two days post infiltration. HR on the spot was noted as being suppressed when less than 50 % of that spot showed cell death following ^77^. Fisher’s exact test was used to analyze the results statistically.

### Callose deposition

Callose deposits were observed according to the protocol with small modifications ^78,79^. Briefly, *NEROSs* or *NEROSs*_*cTPmut*_ or *GFP* (see **Supplementary Table 2** for the vectors used) were infiltrated into *N. benthamiana* leaves using Agrobacterium GV3101 of an OD600 of 0.5. Accordingly, 48 h post infiltration the leaves were treated with 500 nM flg22 for 24 h. After that, they were fixed and destained in a destaining solution (absolute ethyl alcohol: acetic acid, 3:1 v/v) for 24 h followed by 1 h 70 % ethanol, 1 h 50 % ethanol 1 h 10 % NaOH at 37°C. Transparent leaf segments were washed three times in 150 mM K_2_HPO_4_ (pH 9.6) and stained with 0.02 % aniline blue (cat. no: B8563-500ML; Sigma) in 150 mM K_2_HPO_4_ (pH 9.6). The pictures were taken using a wide field epifluorescent microscope (Nikon Instruments Inc.). Images were acquired with DAPI filter (excitation at 387/11 nm and emission 447/60nm). Callose deposits were analyzed in fields of 8.4 mm^2^ using ImageJ ^80^ software and normalized to 1 mm^2^.

### RNA-seq sample preparation

Seeds of tomato (*S. lycopersicum* cv Moneymaker) were surface sterilized in 70% (v/v) ethanol for 10 min and then rinsed three times 5 min with sterile water. The seeds were germinated on MS tissue culture medium containing 4.3 g/l MS medium (Duchefa, catalog no. M0221.0050), 0.5 g/l MES, 30 g/l sucrose, pH 5.8, and 10 g/l agar (Difco, catalog No. 214530) in dark. Seeds were germinated at 24°C growth chamber (16-h-light/8-h-dark photoperiod) for 10-14 days until cotyledons were fully expanded and the true leaves just emerged. *R. rhizogenes* strain ATCC 15834 transformation was performed as described previously by ^81^ with some minor modifications. More specifically, competent rhizogenic Agrobacterium cells were transformed by electroporation ^82^ with the desired binary vector, plated on yeast extract beef (YEB) medium plates with the appropriate antibiotics (100 mg/l spectinomycin), and incubated for 3 to 4 days at 28°C. A transformed Agrobacterium culture was inoculated from fresh plates into YEB liquid medium with the appropriate antibiotics added and grown overnight at 28°C with shaking at 200 rpm. The cut cotyledons were soaked in liquid bacterial suspension at an OD at 600 nm of 0.2-0.4 in MS liquid medium for 20 min and transferred onto MS agar plates without antibiotics for 3 days. After that, the cotyledons were transferred to MS agar plates with 200 mg/l cefotaxime (Duchefa, catalog no. c0111.0025) and 50 mg/l kanamycin and returned to 24°C. The screening for expression of the eGFP marker of antibiotic-resistant roots was monitored by using a wide-field epifluorescent microscope (Nikon Instruments Inc.). Four independent roots showing expression of the marker were subcloned for each construct. After three rounds of cultivation, root cultures were maintained and grown in antibiotic-free MS medium supplemented with 3 % sucrose (w/v) at 24°C. Two weeks after growing in liquid medium effector expression was induced with 100 μM β-Estradiol (cat. no: E8875-1G; Sigma) for 24 h. After that samples were grounded and from one part RNA was extracted, while from the second half proteins were extracted and analyzed by western blot. For mRNA sequencing, RNA was extracted from ±100 mg ground tissue using the RNeasy Plant Mini Kit (Qiagen). Four biological replicates, each independently transformed, were used per treatment. RNA quality control was performed. RNA 6000 nanochip (Agilent technologies) libraries were prepared using QuantSeq 3’ mRNA library prep FWD kit (Lexogen) according to the manufacturer’s instructions. Library-quality was verified using a High sensitivity DNA chip (Agilent technologies). Library quantification was assessed using qPCR assay according to Illumina protocol equimolar pooling of libraries based on qPCR and libraries were sequenced on an Illumina NextSeq 500 platform, SR76, with high output. The sequencing depth was 8M.

### RNA-seq data analysis

Data analyses were carried out using the Galaxy Australia platform (https://usegalaxy.org.au/) ^83^. The quality was assessed using FastQC ^84^ and MultiQC ^85^, and reads were trimmed with Trimmomatic ^86^. Performing initial ILLUMINACLIP step (3,30,10,8) and two additional Trimmomatic steps: SLIDINGWINDOW (3,10) and MINLEN, reads < 20 bp were removed. Trimmed reads were mapped to the tomato reference genome (ITAG 3.0, ^87^) using STAR ^88^. Reads were counted using HTSeq ^89^. Differentially expressed genes were identified using the DeSeq2 ^90^. Genes were considered differentially expressed if P _adjusted_ < 0.01 and log2FC < -1 (downregulated) or log2FC > 1(upregulated). GoSeq ^91^ and g: Profiler ^92^ were used for gene ontology analysis and pathway analysis. Genome and annotation files were assessed from EnsemblPlants (https://plants.ensembl.org/index.html) ^93^.

### RT-qPCR for confirmation of RNA-seq results

RNA was extracted using the RNeasy Plant Mini Kit (Qiagen), and cDNA was synthesized using Maxima First Strand cDNA Synthesis Kit for RT-qPCR (Thermo Fisher Scientific). Primers were designed using QuantPrime ^94^. RT-qPCRs were performed with four biological replicates. The qPCR conditions consisted of initial denaturation at 95 °C for 10 min followed by 50 cycles of 95 °C for 25 s, 58 °C for 25 s, and 72 °C for 20 s (all primers were designed using QuantPrime ^94^ and are listed in the **Supplementary Table 2**. Expression data were normalized using three reference genes ^95^ and analyzed using REST2009 ^96^.

## Acknowledgments

The authors would like to thank Evi Ceulemans and Alain Goossens (Goossens Lab, VIB Center for Plant Systems Biology, Ghent, Belgium) for providing plasmids, strain, and protocol for Y2H and for helpful discussions. Maria Aparicio Chacon and Sofie Goormachtig (Goormachtig Lab, VIB Center for Plant Systems Biology, Ghent, Belgium) for providing the plasmids used for localization studies and helpful discussions. Els Van Damme’s Lab (Faculty of Bioscience Engineering Ghent University, Ghent, Belgium) provided plasmids for markers for localization. Jovana Kaljevic for valuable comments during research design, reading the manuscript, and providing comments and suggestions. The Special Research Fund of Ghent University for the BOF18/GOA/013 project. The visit to INRAE to perform the immunolocalization was funded by FWO (Grant for a short study visit abroad, Number K215922N) and EMBO (Scientific Exchange Grant Application, Number 9604). The grants from the National Natural Science Foundation of China (Number 31171824 and 31471750).

## Author contribution

B.S. and G.G. designed the research; B.S. performed Y2H and callose deposition. B.S. and L.B. performed ROS and HR. B.S. and Y.C. made constructs, *in planta* localization, and prepared samples for RNA-seq. H.X. performed Y2H screening, *in-situ* hybridization, and stage expression analysis. K.S. and H.V.P, B.S., and G.G. designed ETR experiments. B.S. and H.V.P. performed ETR experiments and analyses. B.S. and J.A.E. performed immunolocalization. B.S. performed RNA-seq data analysis, figures, and all the statistical analyses. B.S. and G.G. wrote the manuscript. All authors provided comments.

## Data availability

The data that support the findings of this study are available from the corresponding author upon reasonable request. mRNA-sequencing data are uploaded to Sequence Read Archive (SRA), BioProject ID PRJNA886960.

## References

1. Jones, J. T. et al. Top 10 plant-parasitic nematodes in molecular plant pathology. Mol Plant Pathol 14, 946–61 (2013).

2. Orion, D. & Wergin, W. P. Chloroplast Differentiation in Tomato Root Galls Induced by the Root Knot Nematode Meloidogyne incognita. J Nematol 14, 77–83 (1982).

3. Ji, H. et al. Transcriptional analysis through RNA sequencing of giant cells induced by Meloidogyne graminicola in rice roots. J Exp Bot 64, 3885–3898 (2013).

4. Davis, E. L., Hussey, R. S. & Baum, T. J. Getting to the roots of parasitism by nematodes. Trends Parasitol 20, 134–141 (2004).

5. Huang, G. et al. A profile of putative parasitism genes expressed in the esophageal gland cells of the root-knot nematode Meloidogyne incognita. Mol Plant-microbe Interactions Mpmi 16, 376–81 (2003).

6. Bellafiore, S. et al. Direct Identification of the Meloidogyne incognita Secretome Reveals Proteins with Host Cell Reprogramming Potential. Plos Pathog 4, e1000192 (2008).

7. Siddique, S. & Grundler, F. M. Parasitic nematodes manipulate plant development to establish feeding sites. Curr Opin Microbiol 46, 102–108 (2018).

8. Gheysen, G. & Mitchum, M. G. How nematodes manipulate plant development pathways for infection. Curr Opin Plant Biol 14, 415–421 (2011).

9. Mejias, J., Truong, N. M., Abad, P., Favery, B. & Quentin, M. Plant Proteins and Processes Targeted by Parasitic Nematode Effectors. Front Plant Sci 10, 970 (2019).

10. Grunewald, W. et al. Manipulation of Auxin Transport in Plant Roots during Rhizobium Symbiosis and Nematode Parasitism. Plant Cell 21, 2553–62 (2009).

11. Goverse, A. & Smant, G. The Activation and Suppression of Plant Innate Immunity by Parasitic Nematodes. Annu Rev Phytopathol 52, 243–265 (2014).

12. Song, H. et al. The Meloidogyne graminicola effector MgMO289 targets a novel rice copper metallochaperone to suppress plant immunity. J Exp Bot (2021) doi:10.1093/jxb/erab208.

13. Naalden, D. et al. The Meloidogyne graminicola effector Mg16820 is secreted in the apoplast and cytoplasm to suppress plant host defense responses. Mol Plant Pathol 19, 2416–2430 (2018).

14. Jones, J. D. G. & Dangl, J. L. The plant immune system. Nature 444, 323–329 (2006).

15. Dangl, J. L. & Jones, J. D. G. Plant pathogens and integrated defence responses to infection. Nature 411, 826– 833 (2001).

16. Dodds, P. N. & Rathjen, J. P. Plant immunity: towards an integrated view of plant–pathogen interactions. Nat Rev Genet 11, 539–548 (2010).

17. Toruño, T. Y., Stergiopoulos, I. & Coaker, G. Plant Pathogen Effectors: Cellular Probes Interfering with Plant Defenses in Spatial and Temporal Manners. Annu Rev Phytopathol 54, 1–23 (2015).

18. Thomma, B. P. H. J., Nürnberger, T. & Joosten, M. H. A. J. Of PAMPs and Effectors: The Blurred PTI-ETI Dichotomy. Plant Cell Online 23, 4–15 (2011).

19. Holbein, J., Grundler, F. M. W. & Siddique, S. Plant basal resistance to nematodes: an update. J Exp Bot 67, 2049–2061 (2016).

20. Sato, K., Kadota, Y. & Shirasu, K. Plant Immune Responses to Parasitic Nematodes. Front Plant Sci 10, 1165 (2019).

21. Torres, M. A., Jones, J. D. G. & Dangl, J. L. Reactive Oxygen Species Signaling in Response to Pathogens. Plant Physiol 141, 373–378 (2006).

22. Zabala, M. de T. et al. Chloroplasts play a central role in plant defence and are targeted by pathogen effectors. Nat Plants 1, 15074 (2015).

23. Shapiguzov, A., Vainonen, J. P., Wrzaczek, M. & Kangasjärvi, J. ROS-talk - how the apoplast, the chloroplast, and the nucleus get the message through. Front Plant Sci 3, 292 (2012).

24. Serrano, I., Audran, C. & Rivas, S. Chloroplasts at work during plant innate immunity. J Exp Bot 67, 3845–3854 (2016).

25. Jalmi, S. K. & Sinha, A. K. ROS mediated MAPK signaling in abiotic and biotic stress-striking similarities and differences. Front Plant Sci 6, 769 (2015).

26. Park, E., Nedo, A., Caplan, J. L. & Dinesh-Kumar, S. P. Plant–microbe interactions: organelles and the cytoskeleton in action. New Phytol 217, 1012–1028 (2018).

27. Tripathy, B. C. & Oelmüller, R. Reactive oxygen species generation and signaling in plants. Plant Signal Behav 7, 1621–1633 (2012).

28. Zurbriggen, M. D. et al. Chloroplast-generated reactive oxygen species play a major role in localized cell death during the non-host interaction between tobacco and Xanthomonas campestris pv. vesicatoria. Plant J 60, 962– 973 (2009).

29. Zurbriggen, M. D., Carrillo, N. & Hajirezaei, M.-R. ROS signaling in the hypersensitive response: When, where and what for? Plant Signal Behav 5, 393–396 (2010).

30. Mittler, R., Vanderauwera, S., Gollery, M. & Breusegem, F. V. Reactive oxygen gene network of plants. Trends Plant Sci 9, 490–498 (2004).

31. Xia, X.-J. et al. Interplay between reactive oxygen species and hormones in the control of plant development and stress tolerance. J Exp Bot 66, 2839–2856 (2015).

32. Huang, H., Ullah, F., Zhou, D.-X., Yi, M. & Zhao, Y. Mechanisms of ROS Regulation of Plant Development and Stress Responses. Front Plant Sci 10, 800 (2019).

33. Roldán-Arjona, T. & Ariza, R. R. Repair and tolerance of oxidative DNA damage in plants. Mutat Res Rev Mutat Res 681, 169–179 (2009).

34. Jwa, N.-S. & Hwang, B. K. Convergent Evolution of Pathogen Effectors toward Reactive Oxygen Species Signaling Networks in Plants. Front Plant Sci 08, 1687 (2017).

35. Chopra, D. et al. Plant parasitic cyst nematodes redirect host indole metabolism via NADPH oxidase-mediated ROS to promote infection. New Phytol 232, 318–331 (2021).

36. Xu, Q. et al. An effector protein of the wheat stripe rust fungus targets chloroplasts and suppresses chloroplast function. Nat Commun 10, 5571 (2019).

37. Zhang, J. et al. The DnaJ proteins DJA6 and DJA5 are essential for chloroplast iron–sulfur cluster biogenesis. Embo J e106742 (2021) doi:10.15252/embj.2020106742.

38. Lozano-Durán, R., Bourdais, G., He, S. Y. & Robatzek, S. The bacterial effector HopM1 suppresses PAMP-triggered oxidative burst and stomatal immunity. New Phytol 202, 259–269 (2014).

39. Li, G. et al. Distinct Pseudomonas type-III effectors use a cleavable transit peptide to target chloroplasts. Plant J 77, 310–321 (2014).

40. Lin, B. et al. A novel nematode effector suppresses plant immunity by activating host reactive oxygen species-scavenging system. New Phytologist 209, 1159–73 (2015).

41. Eves-van den Akker, S., Stojilković, B. & Gheysen, G. Recent applications of biotechnological approaches to elucidate the biology of plant–nematode interactions. Curr Opin Biotech 70, 122–130 (2021).

42. Joshi, I. et al. Conferring root-knot nematode resistance via host-delivered RNAi-mediated silencing of four Mi-msp genes in Arabidopsis. Plant Sci 298, 110592 (2020).

43. Maiwald, D. et al. Knock-Out of the Genes Coding for the Rieske Protein and the ATP-Synthase δ-Subunit of Arabidopsis. Effects on Photosynthesis, Thylakoid Protein Composition, and Nuclear Chloroplast Gene Expression. Plant Physiol 133, 191–202 (2003).

44. Jahns, P., Graf, M., Munekage, Y. & Shikanai, T. Single point mutation in the Rieske iron–sulfur subunit of cytochrome b 6/f leads to an altered pH dependence of plastoquinol oxidation in Arabidopsis. Febs Lett 519, 99– 102 (2002).

45. Munekage, Y. et al. Cytochrome b6f mutation specifically affects thermal dissipation of absorbed light energy in Arabidopsis. Plant J 28, 351–359 (2001).

46. Wang, X. et al. Two stripe rust effectors impair wheat resistance by suppressing import of host Fe – S protein into chloroplasts. Plant Physiol 187, 2530–2543 (2021).

47. Danchin, E. G. J. et al. Identification of Novel Target Genes for Safer and More Specific Control of Root-Knot Nematodes from a Pan-Genome Mining. Plos Pathog 9, e1003745 (2013).

48. Rutter, W. B. et al. Mining novel effector proteins from the esophageal gland cells of Meloidogyne incognita. Mol Plant-microbe Interactions Mpmi 27, 965–74 (2014).

49. Grynberg, P. et al. Comparative Genomics Reveals Novel Target Genes towards Specific Control of Plant-Parasitic Nematodes. Genes-basel 11, 1347 (2020).

50. Aspöck, G., Kagoshima, H., Niklaus, G. & Bürglin, T. R. Caenorhabditis elegans Has Scores of hedgehogRelated Genes: Sequence and Expression Analysis. Genome Res 9, 909–923 (1999).

51. Baniulis, D., Yamashita, E., Zhang, H., Hasan, S. S. & Cramer, W. A. Structure–Function of the Cytochrome b6f Complex†. Photochem Photobiol 84, 1349–1358 (2008).

52. Soriano, G. M., Guo, L.-W., Vitry, C. de, Kallas, T. & Cramer, W. A. Electron Transfer from the Rieske Iron-Sulfur Protein (ISP) to Cytochrome f in Vitro IS A GUIDED TRAJECTORY OF THE ISP NECESSARY FOR COMPETENT DOCKING?*. J Biol Chem 277, 41865–41871 (2002).

53. Li, M. & Kim, C. Chloroplast ROS and stress signaling. Plant Commun 3, 100264 (2021).

54. Rodriguez, P. A., Stam, R., Warbroek, T. & Bos, J. I. B. Mp10 and Mp42 from the Aphid SpeciesMyzus persicaeTrigger Plant Defenses inNicotiana benthamianaThrough Different Activities. Mol Plant Microbe In 27, 30– 39 (2014).

55. Renato, M., Boronat, A. & Azcón-Bieto, J. Respiratory processes in non-photosynthetic plastids. Front Plant Sci 6, 496 (2015).

56. Dibrova, D. V., Cherepanov, D. A., Galperin, M. Y., Skulachev, V. P. & Mulkidjanian, A. Y. Evolution of cytochrome bc complexes: From membrane-anchored dehydrogenases of ancient bacteria to triggers of apoptosis in vertebrates. Biochimica Et Biophysica Acta Bba - Bioenergetics 1827, 1407–1427 (2013).

57. Huang, J. et al. An oomycete plant pathogen reprograms host pre-mRNA splicing to subvert immunity. Nat Commun 8, 2051 (2017).

58. Ishikawa, K. et al. Bacterial effector modulation of host E3 ligase activity suppresses PAMP-triggered immunity in rice. Nat Commun 5, 5430 (2014).

59. Zhao, J. et al. The root-knot nematode effector MiPDI1 targets a stress-associated protein, SAP, to establish disease in Solanaceae and Arabidopsis. New Phytologist (2020) doi:10.1111/nph.16745.

60. Moreira, V. J. V. et al. Minc03328 effector gene downregulation severely affects Meloidogyne incognita parasitism in transgenic Arabidopsis thaliana. Planta 255, 44 (2022).

61. Shukla, N. et al. Transcriptome analysis of root-knot nematode (Meloidogyne incognita)-infected tomato (Solanum lycopersicum) roots reveals complex gene expression profiles and metabolic networks of both host and nematode during susceptible and resistance responses. Mol Plant Pathol 19, 615–633 (2018).

62. Coll, N. S., Epple, P. & Dangl, J. L. Programmed cell death in the plant immune system. Cell Death Differ 18, 1247–1256 (2011).

63. Sperschneider, J. et al. LOCALIZER: subcellular localization prediction of both plant and effector proteins in the plant cell. Sci Rep-uk 7, 44598 (2017).

64. Hu, L. et al. Molecular and biochemical characterization of the β-1,4-endoglucanase gene Mj-eng-3 in the root-knot nematode Meloidogyne javanica. Exp Parasitol 135, 15–23 (2013).

65. Jumper, J. et al. Highly accurate protein structure prediction with AlphaFold. Nature 596, 583–589 (2021).

66. Engler, J. de A. et al. Dynamic cytoskeleton rearrangements in giant cells and syncytia of nematode-infected roots. Plant J 38, 12–26 (2004).

67. Kronenberger, J., Desprez, T., Höfte, H., Caboche, M. & Traas, J. A methacrylate embedding procedure developed for immunolocalization on plant tissues is also compatible with in situ hybridization. Cell Biol Int 17, 1013–1021 (1993).

68. Vieira, P. et al. The plant apoplasm is an important recipient compartment for nematode secreted proteins. J Exp Bot 62, 1241–1253 (2010).

69. Pérez, A. C. et al. The Non-JAZ TIFY Protein TIFY8 from Arabidopsis thaliana Is a Transcriptional Repressor. Plos One 9, e84891 (2014).

70. Erffelinck, M.-L. et al. A user-friendly platform for yeast two-hybrid library screening using next generation sequencing. Plos One 13, e0201270 (2018).

71. Kodama*, Y. & Hu, C.-D. An improved bimolecular fluorescence complementation assay with a high signal-to-noise ratio. Biotechniques 49, 793–805 (2010).

72. Bos, J. I. B. et al. A functional genomics approach identifies candidate effectors from the aphid species Myzus persicae (green peach aphid). Plos Genet 6, e1001216 (2010).

73. Felix, G., Duran, J. D., Volko, S. & Boller, T. Plants have a sensitive perception system for the most conserved domain of bacterial flagellin. Plant J 18, 265–276 (1999).

74. Keppler, L. D. Active Oxygen Production During a Bacteria-Induced Hypersensitive Reaction in Tobacco Suspension Cells. Phytopathology 79, 974 (1989).

75. Mendy, B. et al. Arabidopsis leucine-rich repeat receptor-like kinase NILR1 is required for induction of innate immunity to parasitic nematodes. Plos Pathog 13, e1006284 (2017).

76. Kamoun, S., Hamada, W. & Huitema, E. Agrosuppression: A Bioassay for the Hypersensitive Response Suited to High-Throughput Screening. Mol Plant-microbe Interactions 16, 7–13 (2003).

77. Gilroy, E. M. et al. CMPG1-dependent cell death follows perception of diverse pathogen elicitors at the host plasma membrane and is suppressed by Phytophthora infestans RXLR effector AVR3a. New Phytol 190, 653–666 (2011).

78. Schenk, S. & Schikora, A. Staining of Callose Depositions in Root and Leaf Tissues. Bio-protocol 5, (2015).

79. Ji, H., Kyndt, T., He, W., Vanholme, B. & Gheysen, G. β-Aminobutyric Acid–Induced Resistance Against Root-Knot Nematodes in Rice Is Based on Increased Basal Defense. Mol Plant-microbe Interactions 28, 519–533 (2015).

80. Schneider, C. A., Rasband, W. S. & Eliceiri, K. W. NIH Image to ImageJ: 25 years of image analysis. Nat Methods 9, 671–675 (2012).

81. Harvey, J. J. W., Lincoln, J. E. & Gilchrist, D. G. Programmed cell death suppression in transformed plant tissue by tomato cDNAs identified from an Agrobacterium rhizogenes-based functional screen. Mol Genet Genomics 279, 509–521 (2008).

82. Wen-jun, S. & Forde, B. G. Efficient transformation of Agrobacterium spp. by high voltage electroporation. Nucleic Acids Res 17, 8385–8385 (1989).

83. Afgan, E. et al. The Galaxy platform for accessible, reproducible and collaborative biomedical analyses: 2018 update. Nucleic Acids Res 46, W537–W544 (2018).

84. Brandine, G. de S. & Smith, A. D. Falco: high-speed FastQC emulation for quality control of sequencing data. F1000research 8, 1874 (2020).

85. Ewels, P., Magnusson, M., Lundin, S. & Käller, M. MultiQC: summarize analysis results for multiple tools and samples in a single report. Bioinformatics 32, 3047–3048 (2016).

86. Bolger, A. M., Lohse, M. & Usadel, B. Trimmomatic: a flexible trimmer for Illumina sequence data. Bioinformatics 30, 2114–2120 (2014).

87. Hosmani, P. S. et al. An improved de novo assembly and annotation of the tomato reference genome using single-molecule sequencing, Hi-C proximity ligation and optical maps. Biorxiv 767764 (2019) doi:10.1101/767764.

88. Dobin, A. et al. STAR: ultrafast universal RNA-seq aligner. Bioinformatics 29, 15–21 (2013).

89. Anders, S., Pyl, P. T. & Huber, W. HTSeq—a Python framework to work with high-throughput sequencing data. Bioinformatics 31, 166–169 (2015).

90. Love, M. I., Huber, W. & Anders, S. Moderated estimation of fold change and dispersion for RNA-seq data with DESeq2. Genome Biol 15, 550 (2014).

91. Young, M. D., Wakefield, M. J., Smyth, G. K. & Oshlack, A. Gene ontology analysis for RNA-seq: accounting for selection bias. Genome Biol 11, R14–R14 (2010).

92. Raudvere, U. et al. g:Profiler: a web server for functional enrichment analysis and conversions of gene lists (2019 update). Nucleic Acids Res 47, W191–W198 (2019).

93. Bolser, D., Staines, D. M., Pritchard, E. & Kersey, P. Plant Bioinformatics, Methods and Protocols. Methods Mol Biology 1374, 115–140 (2016).

94. Arvidsson, S., Kwasniewski, M., Riaño-Pachón, D. M. & Mueller-Roeber, B. QuantPrime – a flexible tool for reliable high-throughput primer design for quantitative PCR. Bmc Bioinformatics 9, 465 (2008).

95. Cheng, Y. et al. Genome-Wide Identification and Evaluation of Reference Genes for Quantitative RT-PCR Analysis during Tomato Fruit Development. Front Plant Sci 8, 1440 (2017).

96. Pfaffl, M. W. A new mathematical model for relative quantification in real-time RT–PCR. Nucleic Acids Res 29, e45–e45 (2001).

